# Transglutaminase 3 is a Histone Monoaminylation Writer

**DOI:** 10.64898/2026.10.20.760504

**Authors:** Yvonne Ritsema, Huapeng Li, Xingyu Ma, Anne A. Leaman, Qingfei Zheng

## Abstract

Histone monoaminylation, including serotonylation, dopaminylation, and histaminylation, is an emerging class of epigenetic modifications that regulates gene transcription in diverse physiological and disease contexts. Histone H3 glutamine 5 (H3Q5) was the first identified and remains a major site of monoaminylation on core histones. Mechanistically, monoaminylation involves the formation of an isopeptide bond between a substrate glutamine and a monoamine metabolite through a reversible transamidation reaction catalyzed by transglutaminases (TGs). Among the nine members of the human TG family, TG2 has been established as the major writer of H3Q5 monoaminylation. Here, we identify TG3 as an additional histone monoaminylation writer capable of catalyzing H3Q5 histaminylation (H3Q5his).

## Introduction

Histone post-translational modifications (PTMs) constitute a major mechanism for regulating chromatin structure and gene transcription.^1–3^ In addition to canonical PTMs such as methylation, acetylation, phosphorylation, and ubiquitination, an expanding repertoire of metabolite-derived histone modifications has revealed direct biochemical links between cellular metabolism and epigenetic regulation.^4^ Histone monoaminylation represents an emerging class of such modifications in which monoamine metabolites, including serotonin, dopamine, and histamine, are covalently conjugated to glutamine residues on histones.^5^ Histone H3 glutamine 5 (H3Q5) was initially identified as a site of serotonylation (H3Q5ser) and has subsequently emerged as a major site for multiple forms of monoaminylation, including dopaminylation (H3Q5dop) and histaminylation (H3Q5his).^6^ These modifications have been implicated in the regulation of transcription and diverse physiological and pathological processes, highlighting monoamine metabolites as donors for chromatin modification in addition to their established roles in cellular signaling.

Unlike many histone PTMs that are installed through enzymatic transfers from activated donor metabolites (*i*.*e*., co-factors), monoaminylation is generated through a transamidation reaction that directly conjugates the primary amine of a monoamine to the γ-carboxamide group of a target glutamine residue, resulting in formation of a γ-glutamyl monoamine isopeptide bond.^5,6^ This reaction is catalyzed by transglutaminases (TGs), a group of structurally related enzymes that catalyze Ca^2+^-dependent acyl-transfer reactions involving protein glutamine residues.^7^ The key step of this transamidation reaction is the formation of a thioester intermediate between TG’s catalytic cysteine and the target glutamine (**Figure 1A**). Among the nine human TG family members, transglutaminase 2 (TG2) is the most extensively characterized and has been established as the principal writer of histone monoaminylation. Specifically, TG2 catalyzes H3Q5 serotonylation, dopaminylation, and histaminylation and can also mediate the removal or exchange of monoamines through the reversible transamidation reaction.^8^ Despite the presence of multiple catalytically active TG family members with a conserved catalytic architecture (**Figure 1B**), whether histone monoaminylation is uniquely mediated by TG2 or can be written by other transglutaminases remains largely unexplored.

**Figure 1.**
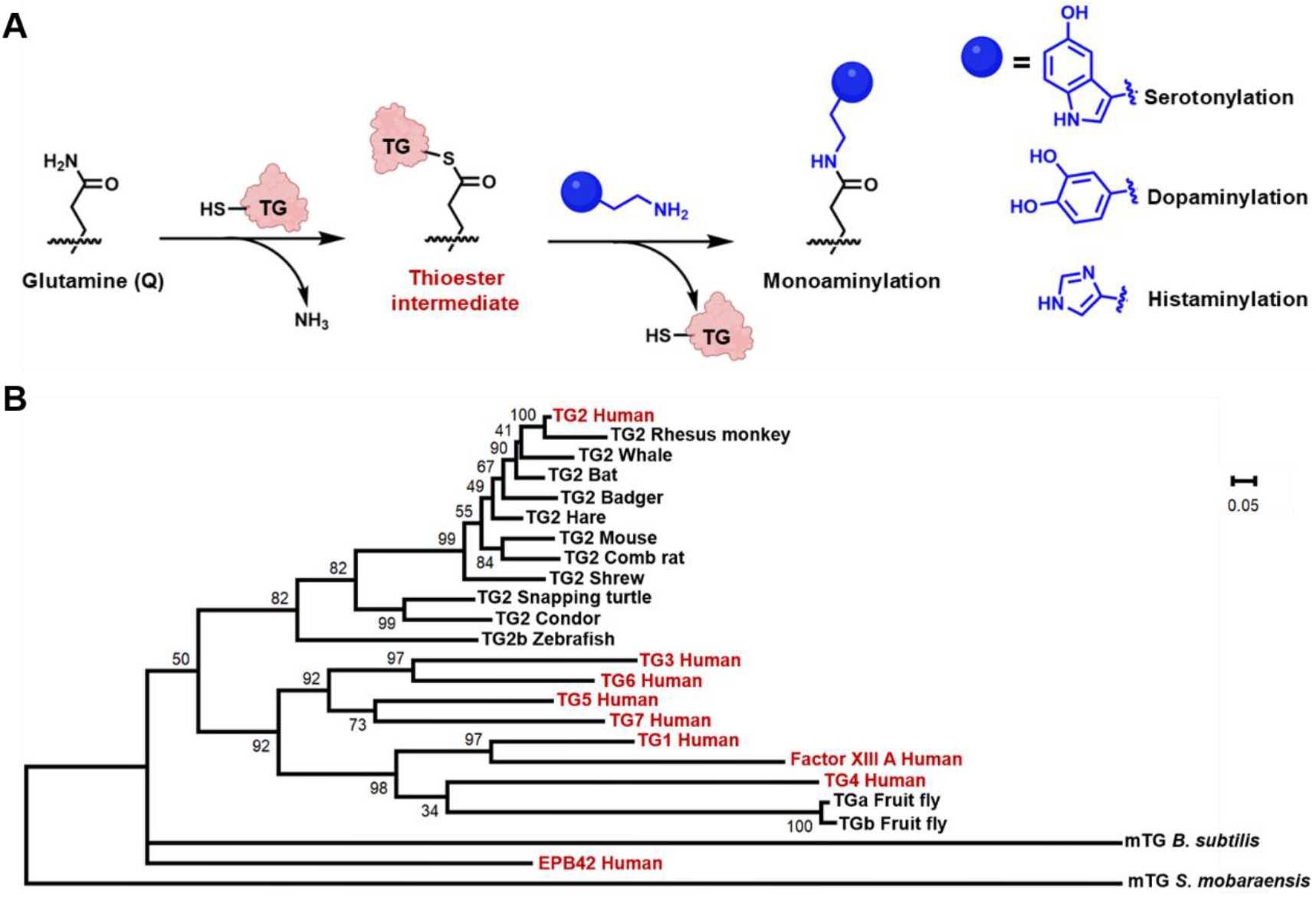
General mechanism of transglutaminase (TG)-catalyzed monoaminylation reaction on substrate glutamines and evolutionary analysis of TGs from different species. (**A**) A catalytic cysteine on TG attacks the γ-carboxamide group of glutamine to produce a thioester intermediate and release ammonia. The thioester intermediate undergoes an aminolysis reaction with a primary amine on various monoamine donors to release TG and covalently conjugate the monoamine donors (*e*.*g*., serotonin, dopamine, and histamine) to glutamine. (**B**) Phylogenetic analysis of TGs from different species. The tree was constructed with the Neighbor-Joining (NJ) method. Nine human TGs were labeled in red. Scale bar, 0.05 substitutions per site.

Transglutaminase 3 (TG3) is a Ca^2+^-dependent transamidase best known for its functions in epithelial differentiation and protein cross-linking.^9^ TG3 shares the conserved catalytic machinery of the transglutaminase family with TG2 but exhibits distinct patterns of expression, activation, and substrate recognition (**Figure 2**).^10–12^ Whether TG3 can utilize monoamine metabolites as amine donors and catalyze histone monoaminylation has not been established. Here, using *in vitro* biochemical reconstitution and cellular analyses, we identify TG3 as an additional writer of histone monoaminylation. We demonstrate that TG3 efficiently utilizes histamine as an amine donor and catalyzes histaminylation of histone H3 at Q5, establishing H3Q5his as a product of TG3-mediated transamidation. These findings expand the enzymatic repertoire underlying histone monoaminylation beyond TG2 and reveal TG3 as a previously unrecognized writer of H3Q5his.

**Figure 2.**
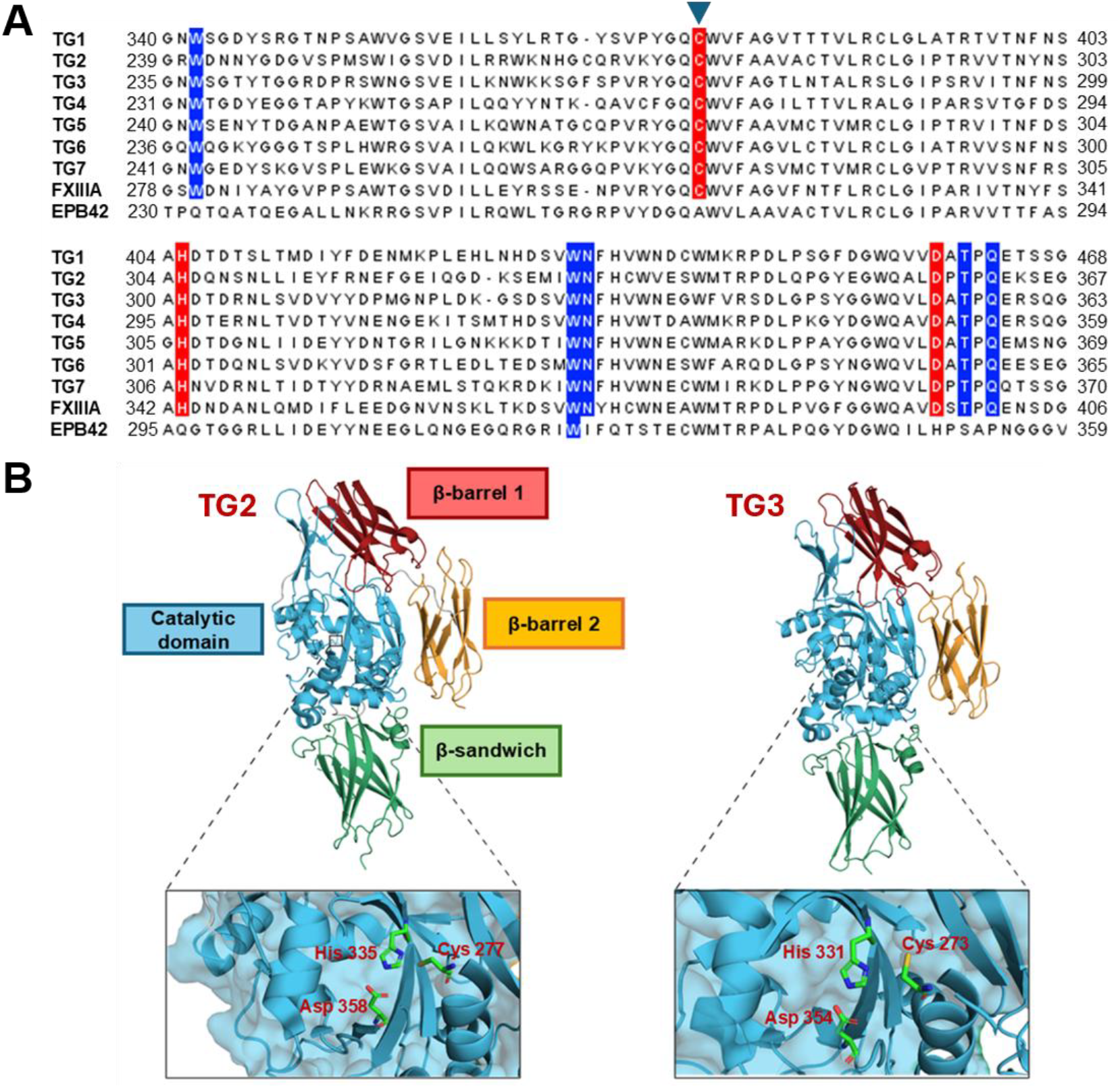
(A) The protein sequences for all nine members of the human TG family were aligned. Shown here is a portion of the catalytic domain with the catalytic triad highlighted in red and other key catalytic and stabilizing residues highlighted in blue. The catalytic cysteines of these human TGs were indicated using an inverted triangle. (**B**) Three-dimensional structure comparison between TG2 (PDB: 6KZB) and TG3 (PDB: 1NUD) highlighting the four primary structural domains and conserved catalytic triad.

## Results and Discussion

To identify additional potential writers of histone monoaminylation, we performed comparative bioinformatic analyses of transglutaminases (TGs) from humans and other species (**Figures 1B** and **2A**). Structural comparison revealed that TG2 and TG3 share a similar overall architecture and a conserved catalytic triad (**Figure 2B**). However, unlike TG2, full-length TG3 is translated as a zymogen that requires proteolytic cleavage of a flexible loop for activation (**Figure S1**).^10–12^ To investigate the histone monoaminylation activities of different TG family members, we recombinantly expressed and purified His6-tagged human TG1-TG6 using immobilized metal affinity chromatography (**Figure 3A**). Notably, none of these enzymes were subjected to proteolytic activation before the subsequent *in vitro* biochemical assays. Interestingly, spontaneous cleavage of TG3 was observed by using sodium dodecyl sulfate-polyacrylamide gel electrophoresis (SDS-PAGE) analysis (**Figure S2**). High-resolution protein mass spectrometry further revealed that the combined molecular masses of the N- and C-terminal fragments closely matched that of full-length TG3, suggesting the existence of a single cleavage site during this self-activation process (**Figure S3**).

**Figure 3.**
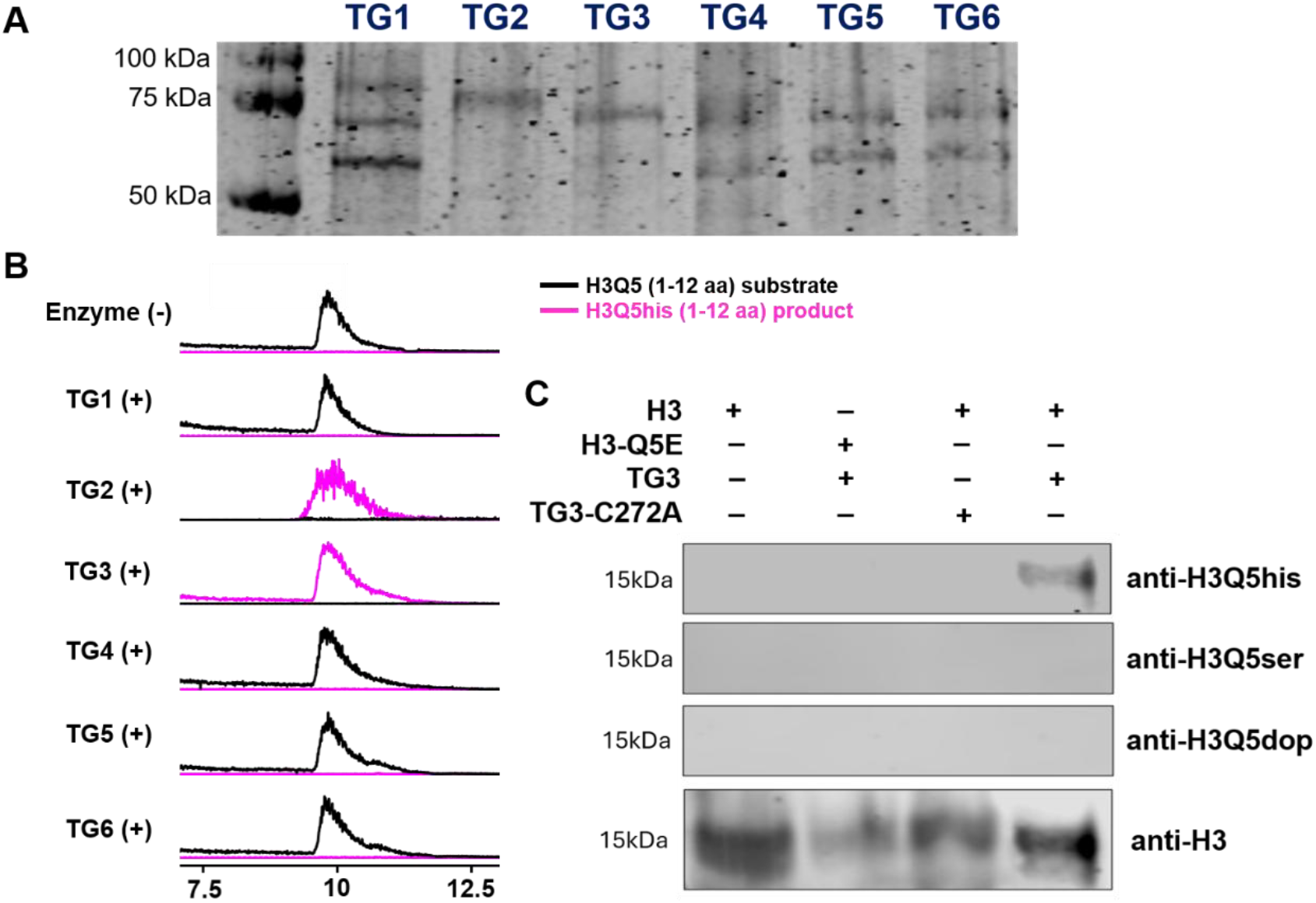
Expression and purification of recombinant human TGs 1-6 and *in vitro* enzymatic activity of TG3 for catalyzing H3Q5his using a peptide or full-length H3 as substrates. (**A**) SDS-PAGE analysis of purified human TGs 1-6, indicating the spontaneous cleavages of TGs 1, 3, 4, 5, and 6. (**B**) HPLC-MS analysis of the conversion from H3Q5 (1-12 aa) substrate peptide to H3Q5his (1-12 aa) catalyzed by TG2 and TG3 in the presence of histamine. TGs 1, 4, 5, and 6 did not show any enzymatic activity on the peptide substrate under the same conditions. (**C**) Immunoblotting analysis of recombinant full-length histone H3 variants (wild-type and Q5E) treated with TG3 or TG3-C272A in the presence of corresponding monoamines (histamine, serotine, or dopamine). Site-specific antibodies recognizing each PTM were used to illustrate the *in vitro* biochemical reactions.

Using a synthetic peptide corresponding to the N-terminal 12 residues of histone H3 as the substrate,^13^ we screened the purified TGs for histone monoaminylation activity. Liquid chromatography-mass spectrometry (LC-MS) analysis revealed that, among the six TGs examined, only TG2 and TG3 catalyzed detectable monoaminylation of the H3 peptide under these assay conditions (**Figures 3B** and **S4**). Subsequent LC-MS-based kinetic analyses demonstrated that TG3 exhibited a pronounced preference for histamine over dopamine and serotonin as a monoamine donor (**Figure S5**).^14^ To further characterize the histone monoaminylation activity of TG3, we generated and purified a catalytically inactive mutant, TG3-C272A. Full-length histone H3 proteins, including wild-type H3 and the H3Q5E mutant,^15^ were also purified as substrates. In the presence of Ca^2+^ and individual monoamine donors (serotonin, dopamine, or histamine), the H3 substrates were incubated with either wild-type TG3 or TG3-C272A. Immunoblotting analyses using modification-specific antibodies revealed that wild-type TG3 efficiently catalyzed the formation of H3Q5his,^16^ whereas the catalytically inactive TG3-C272A mutant failed to generate detectable H3Q5his (**Figure 3C**). In contrast to TG2, TG3 did not produce detectable H3Q5 serotonylation or dopaminylation under the same *in vitro* assay conditions (**Figure 3C**). Further pH-dependence analysis demonstrated that TG3-catalyzed H3Q5his strongly favors more alkaline conditions *in vitro* (**Figure S6**). Collectively, these findings establish TG3 as an additional writer enzyme of the epigenetic mark H3Q5his and suggest that TG3 exhibits more restricted monoamine donor specificity than TG2.

Finally, to investigate whether TG3 catalyzes H3Q5his in cells, we performed ectopically expression of wild-type TG3 or the catalytically inactive TG3-C272A mutant in HEK 293T cells supplemented with histamine. Notably, endogenous TG2 or TG3 expression was undetectable in HEK 293T cells under experimental conditions, providing a suitable cellular platform for evaluating their respective enzymatic activities.^8,13–16^ Histones were extracted at different time points following transfection and analyzed by immunoblotting using an H3Q5his-specific antibody. Expression of wild-type TG3 resulted in a time-dependent increase in H3Q5his levels, with higher histaminylation detected at day 3 than at day 1, whereas TG3-C272A failed to induce any detectable H3Q5his formation (**Figure 4A**). In contrast, ectopic expression of TG2 as a positive control produced a different temporal pattern. H3Q5his levels were substantially elevated on day 1 but declined to undetectable levels by day 3 (**Figure 4A**). This observation is consistent with the previously established reversible transamidation activity of TG2, which enables the installation, removal, and exchange of histone monoaminylation.^8^ In contrast, TG3 exhibited no detectable eraser or monoamine-exchange activity under the *in vitro* or cellular conditions examined, suggesting distinct catalytic properties of these two histone monoaminylation writers. Furthermore, time-dependent cell viability analysis revealed that the overexpression of wild-type TG3 in histamine-treated 293T cells could promote cell growth, similarly to TG2 overexpression (**Figure 4B**). Collectively, these biochemical and cellular findings identify TG3 as an additional writer of H3Q5his and reveal differences between TG2 and TG3 in their catalytic selectivity and regulation of this novel histone PTM, expanding our understandings of the enzymology of this emerging epigenetic mark.

**Figure 4.**
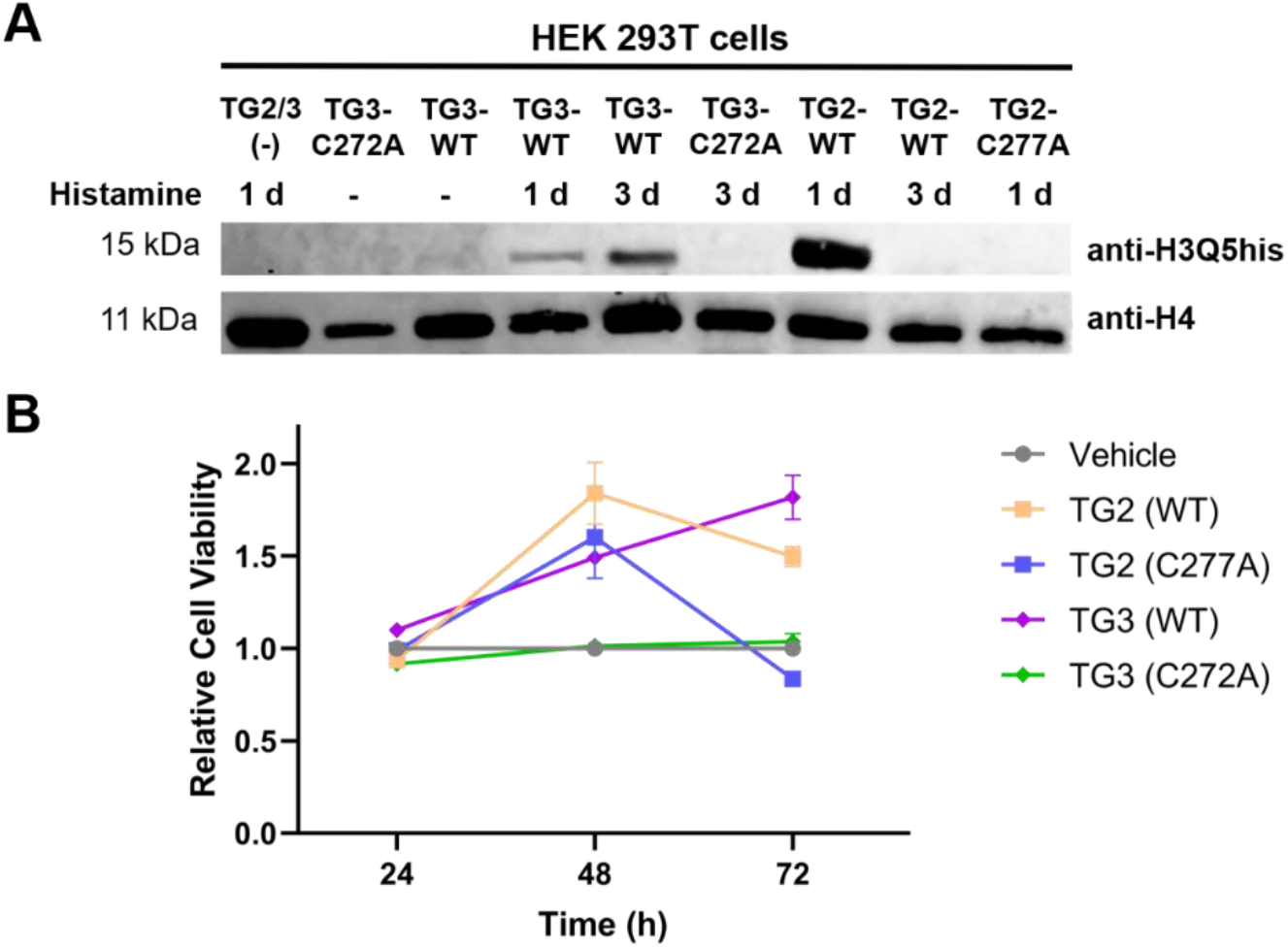
Expression of TG3 and its catalytically inactive mutant (TG3-C272A) in HEK 293T cells, showing its biochemical activity inducing H3Q5his *in cellulo* and impacts on cell viability. (**A**) Immunoblotting analysis of H3Q5his formation catalyzed by TG2 or TG3. The TG2, TG2-C277A, TG3, and TG3-C272A plasmids were transfected into HEK 293T cells with or without histamine treatment. The cells were lysed and histone fractions were extracted after one day (1 d) or three days (3 d). Anti-H3Q5his was used for illustrating the PTM in cells and anti-H4 was used as the loading control. (**B**) Cell viability of TG2, TG2-C277A, TG3, and TG3-C272A-overexpressed 293T cells treated with histamine. Error bars represent the standard deviation from three different experiments. Data are presented as mean values ± SEM

## Conclusion

TGs are a family of cysteine-dependent multifunctional enzymes that catalyze protein transamidation through the formation of a thioester intermediate between the catalytic cysteine residue and γ-carboxamide group of a substrate glutamine (**Figure 1A**).^7,12^ Among the nine members of the human TG family, TG2 is the most widely expressed and extensively studied. TG2-mediated histone monoaminylation has emerged as an important class of epigenetic modifications involved in transcriptional regulation. H3Q5 was the first identified monoaminylation site on core histones and has since attracted considerable research interest.^5,6^ Recently, our laboratory demonstrated that histamine is a preferred monoamine donor for TG2-mediated H3Q5 monoaminylation and identified seven additional glutamine residues on other core histones (including H2AX, H2B, and H4) undergoing histaminylation.^14^ Despite these advances, whether other human TGs can function as histone monoaminylation writers remains largely unexplored.

In this study, we recombinantly expressed and purified six human TG family members (TG1-TG6) and evaluated their monoaminylation activities using an H3-derived peptide substrate. LC-MS and site-specific antibody-based immunoblotting analyses identified TG3 as an additional writer enzyme capable of catalyzing H3Q5his. In contrast to TG2, TG3 exhibited a pronounced preference for histamine, whereas serotonylation and dopaminylation of full-length H3 were not detectable under the assay conditions examined. Notably, unlike conventional TG3 activation protocols, our biochemical assays did not involve any exogenous proteolytic treatment with dispase or other proteases.^10,11^ Instead, spontaneous cleavage of recombinant TG3 was observed, with LC-MS analysis consistent with a single self-cleavage site. Subsequent cellular experiments demonstrated that ectopic expression of TG3 induced H3Q5his in HEK 293T cells, although the modification accumulated more slowly than in TG2-expressing cells. Furthermore, TG3 exhibited no detectable H3Q5his removal or monoamine-exchange activity under the conditions examined, suggesting distinct catalytic properties of these two histone monoaminylation writers.^8^ In summary, these biochemical and cellular findings establish TG3 as an additional writer of H3Q5his, expanding the enzymatic mechanisms of histone monoaminylation^17^ and providing new insights into the biochemical regulation of metabolite-derived histone modifications.^18–20^

## Data availability

The data supporting this article have been included as part of the ESI available at [https://doi.org/XXXXXX].

## Acknowledgements

This research work was financially supported by the NIH (R35 GM150676) and startup funds from Purdue University for Q. Z.

## Accession IDs

TG1: P22735; TG2: P21980; TG3: Q08188; TG4: P49221; TG5: O43548; TG6: O95932; H3: Q71DI3

## Visual Abstract

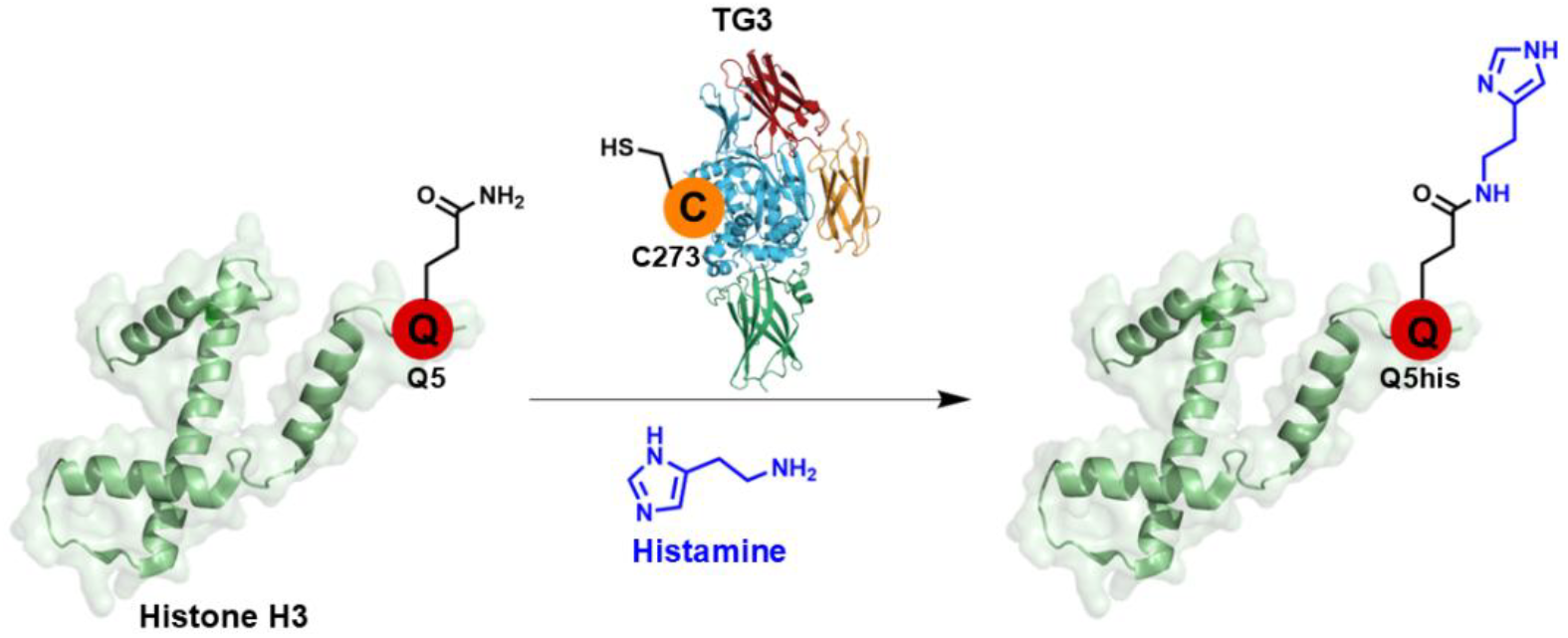

