## Supplemental Information for "Transglutaminase 3 is a Histone Monoaminylation Writer"

### Materials and Methods

#### General methods

UV spectrometry was performed on a NanoDrop 2000c (Thermo Scientific). Biochemicals and media were purchased from Fisher Scientific or Sigma-Aldrich Corporation unless otherwise stated. T4 DNA ligase, DNA polymerase and restriction enzymes were obtained from New England BioLabs. PCR amplifications were performed on an Applied Biosystems Veriti Thermal Cycler using either Taq DNA polymerase (Vazyme Biotech) for routine genotype verification or Phanta Max Super Fidelity DNA Polymerase (Vazyme Biotech) for high-fidelity amplification. Site specific mutagenesis was performed according to standard procedures of the QuickChange Site-Directed Mutagenesis Kit purchased from Stratagene (GE Healthcare) or Mut Express II (Vazyme Biotech). Primer synthesis and DNA sequencing were performed by Integrated DNA Technologies and Genewiz, respectively. PCR amplifications were performed on a Bio-Rad T100TM Thermal Cycler. Centrifugal filtration units were purchased from Millipore, and MINI dialysis units purchased from Pierce. Size exclusion chromatography was performed on an AKTA FPLC system from GE Healthcare equipped with a P-920 pump and UPC-900 monitor. Sephacryl S-200 columns were obtained from GE Healthcare. The primers used in this study for cloning were listed in **Supplementary Table 1**. All the western blots were performed using the primary antibodies annotated in **Supplementary Table 2** and fluorophore-labeled secondary antibodies annotated in **Supplementary Table 3** following protocols recommended by the manufacture. Blots and Cy5-azid gels were imaged on an Odyssey CLx Imaging System (Li-Cor). Additional stained gels were imaged on BioRad Geldck Imager.

#### Peptide synthesis

Standard Fmoc-based Solid Phase Peptide Synthesis (FmocSPPS) was used for the synthesis of peptides in this study. Generally, the peptides were synthesized on ChemMatrix resins with Rink Amide to generate C-terminal amides. Peptides were synthesized using manual addition of the reagents (using a stream of dry $N_2$ to agitate the reaction mixture). For amino acid coupling, 5 eq. Fmoc protected amino acid were pre-activated with 4.9 eq. HBTU, 5 eq. HOBt and 10 eq. DIPEA in DMF and then reacted with the N-terminally deprotected peptidyl resin. Fmoc deprotection was performed in an excess of 20% (v/v) piperidine in DMF, and the deprotected peptidyl resin was washed thoroughly with DMF to remove trace piperidine. Cleavage from the resin and side-chain deprotection were performed with 95 % TFA, 2.5% TIS and 2.5% $H_2O$ at room temperature for 1.5 hours. The peptides were then precipitated with cold diethyl ether, isolated by centrifugation and dissolved in water with 0.1 % TFA followed by RP-HPLC and ESI-MS analyses. Preparative RP-HPLC was used to purify the peptides of interest.[1-3]

#### Expression of recombinant WT TGs 1, 3, 4, 5, 6 and TG3-C272A mutant

The W118-1_hTG-Flag plasmids for TG1, 3-6 were a gift from Huda Zoghbi (Addgene plasmid # 180406, #180408-180411).[4] First, a ~2000-bp DNA fragment comprising the *tg* gene insert was

PCR amplified using primers corresponding to each TG (1, 3-6) (Supplementary Table 1) and cloned into pet-28a(+) backbone containing a C-terminal His tag using Gibson Assembly, creating pet-28a(+)_TG1/TG3-6. The TG3 C272A mutation was cloned by site-directed mutagenesis using pet-28a(+)_TG3 or W118-1_hTG3-Flag as the template and the primers G160 and G161 (Table 2).

**Purification of recombinant wild-type TGs 1-6 and TG3-C272A mutant**

The wild-type TG1-6 and mutant TG3 $His_8$-tagged proteins were expressed in E. coli BL21 (DE3) cells with an overnight IPTG induction at 16 °C. The bacterial pellet was lysed by sonication, and the lysate was cleared by centrifugation at 11,000 r.p.m. for 60 minutes. The lysate was loaded on Ni-NTA resin and eluted stepwise with elution buffer 50 mM Tris HCl, 300 mM NaCl, 300 mM imidazole, pH 8.0), followed by desalting using PD midiTrap G-25 sephadex columns according to the manufacturer's protocol. Purified recombinant proteins were analyzed by SDS-PAGE and concentrated using stirred ultrafiltration cells (Millipore) according to the manufacturer's protocol. The concentration of each protein was determined using 280 nm wavelength on a NanoDrop 2000c (Thermo Scientific).

**Recombinant histone expression and purification**

Recombinant human histones H3(wild-type and H3Q5E mutant) were expressed in *E. coli* BL21 (DE3) or *E. coli* C41 (DE3), extracted by guanidine hydrochloride and purified by flash reverse chromatography, as previously described.[1] The purified histones were analyzed by ZenoTof: H3.2 $[M + H]^+$ observed, 15256.95 Da; expected, 15256.8 Da.

***In vitro* biochemical assays**

For peptide labeling assays, reactions were carried out in a total volume of 20 μL containing Tris-HCl buffer (100 mM, pH 8.0), $CaCl_2$ (5.0 mM), histamine (10 mM), H3 (1-12 aa) peptide (1 mM) and TG3-WT/TG3-C272A (4 μM). The reaction mixtures were incubated at 37 °C overnight and quenched by the addition of acetonitrile (20 μL). The samples were centrifuged to remove precipitated proteins, and the supernatants were subjected to LC-MS analysis. For protein labeling assays, reactions were carried out in a total volume of 30 μL containing Tris-HCl buffer (100 mM, pH 8.0), $CaCl_2$ (5.0 mM), histamine, serotonin, or dopamine (10 mM), histone H3 protein (1 mM) and TG3-WT/TG3-C272A (4 μM). The reaction mixtures were incubated at 37 °C for 3h and subjected to were analyzed using SDS-PAGE and western blot. Briefly, proteins were analyzed using 12% SDS-PAGE gels, which were transferred from the gel to a PVDF membrane (Life Technologies) using a membrane transfer system (Bio-Rad). The membrane was blocked with 5% milk for 2 h at room temperature with gentle swinging and incubated with H3Q5his, H3Q5ser, or H3Q5dop primary antibody at 4 °C overnight after 4 time washing (5 minutes per time). On the next day, antibody solution was removed, and membrane was incubated with and the secondary antibody (1:10000) at room temperature for 1 hour. The membrane was washed 4 times before signal detection by the imaging system (ODYSSEY CLX). The membrane was also incubated with

H3 primary antibody at 4 °C overnight and then incubated with the secondary antibody (1:10000) at room temperature for 1 hour before imaging.

**Kinetic analysis of TG3-catalyzed transamidation**

To determine the kinetic parameters of TG3-catalyzed transamidation in the presence of different monoamine substrates, the reactions were performed at 30 °C in a final volume of 33.18 µL containing Tris-HCl buffer (pH 8.0), 5 mM $CaCl_2$, 0.25 mM H3 (1-12 aa) peptide, 5.275 µM recombinant TG3, and varying concentrations of monoamine substrates. Histamine, serotonin, and dopamine were tested as monoamine substrates at final concentrations of 0, 0.1, 0.25, 0.50, 1, 2, 4, 6, and 10 mM. After incubation for 3 h, the reactions were quenched with 33.18 µL ACN and analyzed by LC-MS. The formation of monoamine-modified peptide products was quantified based on extracted ion chromatogram peak areas. The resulting velocities were fitted to the Michaelis-Menten equation using Graphpad Prism 9.5 to obtain $K_m$ and $k_{cat}$ values.

**pH-Dependence analysis of TG3-catalyzed histaminylation**

To determine the pH tolerance of TG3-catalyzed histaminylation in the presence of different pH conditions, the reactions were performed at 30 °C in a final volume of 30 µL containing 5 mM $CaCl_2$, 10 mM histamine, 0.25 mM H3 (1-12 aa) peptide, 5.275 µM recombinant TG3 or TG3-C272A, and varying buffer conditions. pH was tested with 100 mM MES pH 5.0, 100 mM MES pH 6.0, 100 mM HEPES pH 7.0, 100 mM Tris-HCl pH 8.0, and 100 mM Tris-HCl pH 9.0. After incubation for 3 h, the reactions were quenched with 30 µL ACN and analyzed by LC-MS. The formation of monoamine-modified peptide products was quantified based on extracted ion chromatogram peak areas.

**Expression of Flag-tagged TG2 and TG3 in HEK 293T cells**

The plasmid of TG2 and its mutant were constructed expressed in HEK 293T cells based on the protocol of our previous research.[2] Briefly, wild-type TG2, wild-type TG3, the TG2-C277A mutant, and the TG3-C272A mutant were overexpressed in HEK 293T cells using Lipofectamine 2000 Transfection Reagent (Thermo Fisher Scientific) according to the manufacturer's protocol. HEK 293T cells (ATCC) were cultured at 37 °C with 5% CO2 in DMEM medium supplemented with 10% fetal bovine serum (FBS) (Sigma-Aldrich), 2 mM Lglutamine and 500 units mL−1 penicillin and streptomycin.

**Identification of H3 glutamine 5 histaminylation (H3Q5his) from cells**

HEK293T cells transfected with WT TG2, TG2-C277A mutant, TG3, or TG3-C272A mutant plasmids were treated with 5 mM histamine for 24 or 72 hours. Briefly, the cell lysis solution was prepared using hypotonic lysis buffer (10 mM Tris-HCl pH 8.0, 1 mM KCl, 1.5 mM $MgCl_2$ and 1 mM TCEP, protease and phosphatase inhibitors to 1 × from stock). Histones were then extracted from collected cell pellets, as described in our previous research[5] for further western blot analysis. Briefly, the chromatin pellet was resuspended in 0.4 N $H_2SO_4$ and rotated at 4 °C overnight to extract histones. After centrifugation at 16,000 X g for 10 minutes, the supernatant was transferred to a new tube followed by adding 132 μL of 100% TCA to precipitate histones for 1 hour. At last, histones were dried at room temperature after precipitation by centrifugation. The dried histones were then dissolved in 25 μL 4X SDS loading buffer and were analyzed using SDS-PAGE and western blot. Briefly, proteins were analyzed using 12% SDS-PAGE gels, which were transferred from the gel to a PVDF membrane (Life Technologies) using a membrane transfer system (Bio-Rad). The membrane was blocked with 5% milk for 2 h at room temperature with gentle swinging and incubated with H3Q5his first antibody at 4 °C overnight after 4 time washing (5 minutes per time). On the next day, antibody solution was removed, and membrane was incubated with and the secondary antibody (1:10000) at room temperature for 1 hour. The membrane was washed 4 times before signal detection by the imaging system (ODYSSEY CLX). The membrane was also incubated with H4 primary antibody at 4 °C overnight and then incubated with the secondary antibody (1:10000) at room temperature for 1 hour before imaging.

**CCK-8 assay for cell viability**

TG2-WT, TG2-C277A, TG3-WT, and TG3-C272A expressing HEK 293T cells were seeded in 96-well plates and cultured under standard conditions. Cell viability was evaluated at 24, 48, and 72 h using a Cell Counting Kit-8 (CCK-8) assay according to the manufacturer's instructions (MCE Cat No HY-K0301). At each time point, CCK-8 solution was added to each well, followed by incubation at 37 °C for 1 h. The absorbance was measured at 450 nm using a microplate reader. Relative cell viability was calculated by normalizing the absorbance values of TG2-WT, TG2-C277A, TG3-WT, and TG3-C272A cells to those of HEK 293T cells treated with vehicle at the same time point, with the vehicle treated cells set to 1.0.

**Evolutionary analysis of TGs**

The bootstrap consensus tree inferred from 1,000 replicates is taken to represent the evolutionary history of the taxa analyzed where branches corresponding to partitions reproduced in less than 50% of replicate trees are collapsed. The percentage of replicate trees in which the associated taxa clustered together (1,000 replicates) is shown next to the branches. The initial tree for the heuristic search was selected by choosing the tree with the superior log-likelihood between a Neighbor-Joining (NJ) tree and a Maximum Parsimony (MP) tree. The NJ tree was generated using a matrix of pairwise distances computed using the Le-Gascuel (2008) LG model. The MP tree had the shortest length among 10 MP tree searches, each performed with a randomly generated starting tree. The analytical procedure encompassed 24 amino acid sequences with 944 positions in the final dataset. Evolutionary analyses were conducted in MEGA12 utilizing up to 4 parallel computing threads.[6]

**Statistics and reproducibility**

For nucleosomal array compaction and cell survival assays, data are presented as the mean ± S.E.M. of three independent experiments. All the western blotting and mass spectrometry data were repeated independently three times with similar results. Statistical analyses were performed in Microsoft Excel, GraphPad Prism 7 with ANOVA, the Student's *t*-test, or $\chi^2$ test.

#### Supplementary Tables

**Supplementary Table 1.** Primers used for molecular cloning in this study.

| Primer Name | Sequence (5' to 3') |
|---|---|
| YR21_pet28a_TG1_F | ggcatctcgaggtggagctcaccaccaccaccaccactgagatccgg |
| YR22_TG1_R | ccggatctcagtggtggtggtggtggtgagctccacctcgagatgcc |
| YR23_TG1_F | aagaaggagatataccatgatggatgggccacgttccgatgtgggccg |
| YR24_pet28a_TG1_R | cggcccacatcggaacgtggcccatccatcatggtatatctccttctt |
| YR25_TG3_F | agaaggagatataccatggctgctctaggagtccagagtatcaactgg |
| YR26_pet28a_TG3_R | ccagttgatactctggactcctagagcagccatggtatatctccttct |
| YR27_pet28a_TG3_F | gatgtggccgaacaccaccaccaccaccactgag |
| YR28_TG3_R | ctcagtggtggtggtggtggtgttcggccacatc |
| YR29_TG4_F | aagaaggagatataccatgatggatgcatcaaaagagctgcaagttc |
| YR30_pet28a_TG4_R | gaacttgcagctcttttgatgcatccatcatggtatatctccttctt |
| YR31_pet28a_TG4_F | gttctcatcaccaagcaccaccaccaccacc |
| YR32_TG4_R | ggtggtggtggtggtgcttggtgatgagaac |
| YR33_TG5_F | aagaaggagatataccatggcccaagggctagaagtgg |
| YR34_pet28a_TG5_R | ccacttctagcccttgggccatggtatatctccttctt |
| YR35_pet28a_TG5_F | gtttatgtagactttgcattacaccaccaccaccaccactga |
| YR36_TG5_R | tcagtggtggtggtggtggtgtaatgcaaagtctacataaac |
| YR37_TG6_F | agaaggagatataccatggcagggatcagagtcaccaaggtggactggc |
| YR38_pet28a_TG6_R | gccagtccaccttggtgactctgatccctgccatggtatatctccttct |
| YR39_pet28a_TG6_F | gtggccactgccaagcaccaccaccaccacca |
| YR40_TG6_R | tggtggtggtggtggtgcttggcagtggccac |
| HLG160_TG3_C272A_F | gatatggccaggcctgggtctttgctggg |
| HLG161_TG3_C272A_R | cccagcaaagacccaggcctggccatatc |

**Supplementary Table 2.** Primary antibodies used in this study.

| Host | Epitope | WB | Vendor |
|---|---|---|---|
| Rabbit | Anti-H3Q5his | 1:1000 | MilliporeSigma (ABE2578) |
| Rabbit | Anti-H3Q5ser | 1:1000 | MilliporeSigma (ABE1791) |
| Rabbit | Anti-H3Q5dopaminyl | 1:1000 | MilliporeSigma (ABE2588) |
| Mouse | Anti-H3 | 1:1000 | CST (1B1B2) |
| Rabbit | Anti-H4 | 1:1000 | Abcam (AB10158) |

**Supplementary Table 3.** Secondary antibodies used in this study.

| Host | Epitope | Label | Dilution | Vendor |
|---|---|---|---|---|
| Donkey | Anti-rabbit | IRDye 680RD | 1:15000 | Li-Cor |
| Goat | Anti-mouse | IRDye 800CW | 1:15000 | Li-Cor |

#### Supplementary Figures

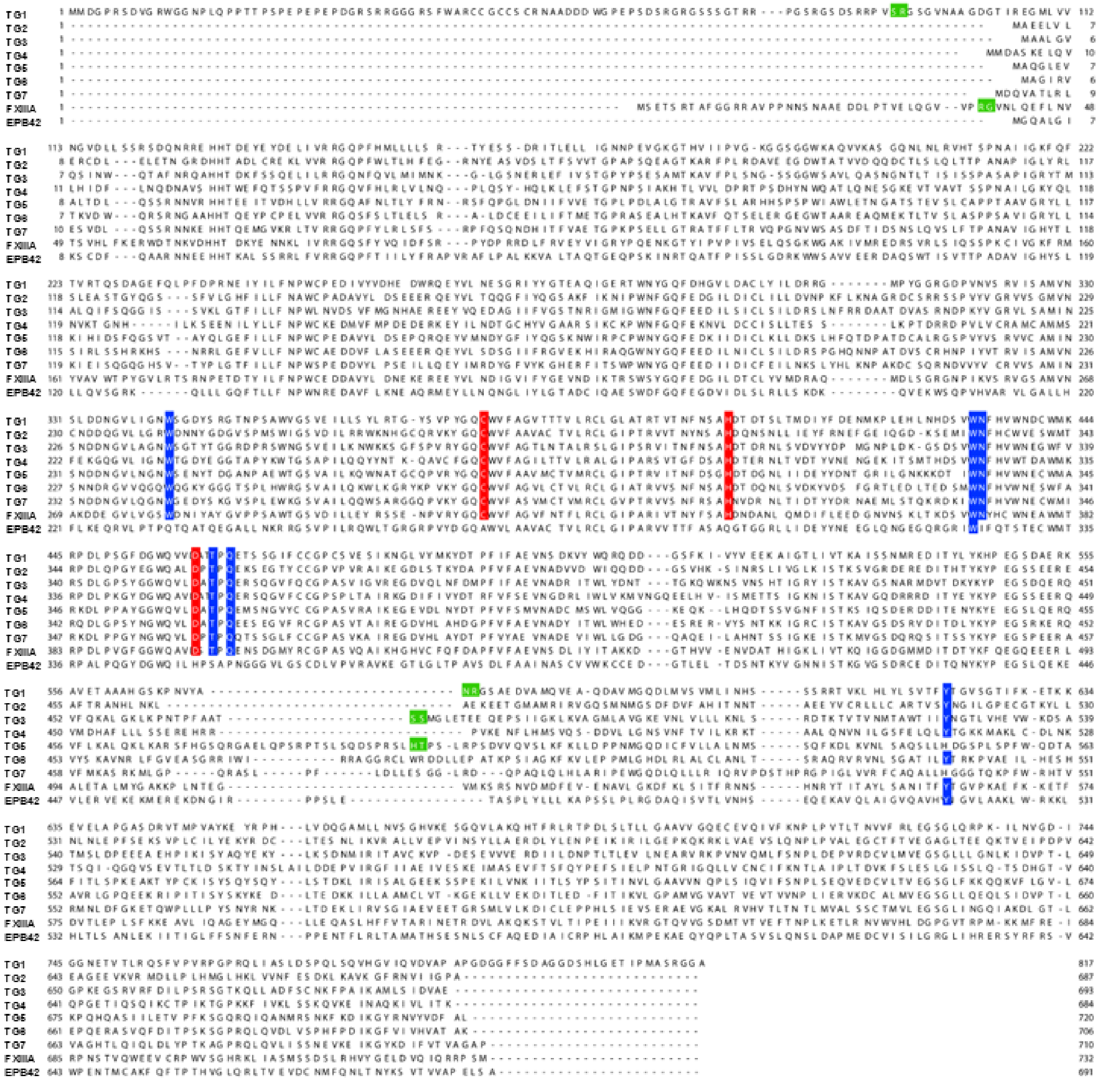

**Figure S1.** Full protein sequence alignment for all nine members of the human transglutaminase family. Residues highlighted in red are the conserved catalytic triad, residues highlighted in blue are other key catalytic and stabilizing residues, and residues highlighted in green are previously reported zymogen cleavage sites.

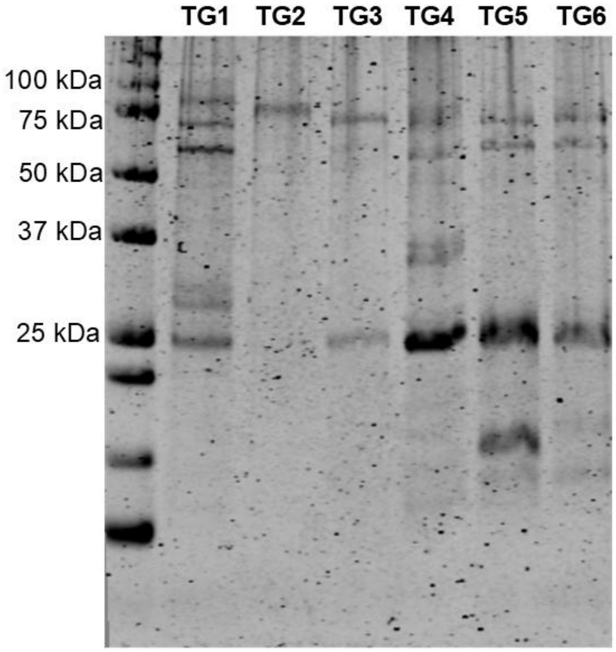


**Figure S2**. Full SDS-PAGE gel analysis of recombinantly purified His$_6$-tagged human TGs 1-6, indicating the self-cleavage and activation of TGs 1, 3, 4, 5, and 6.

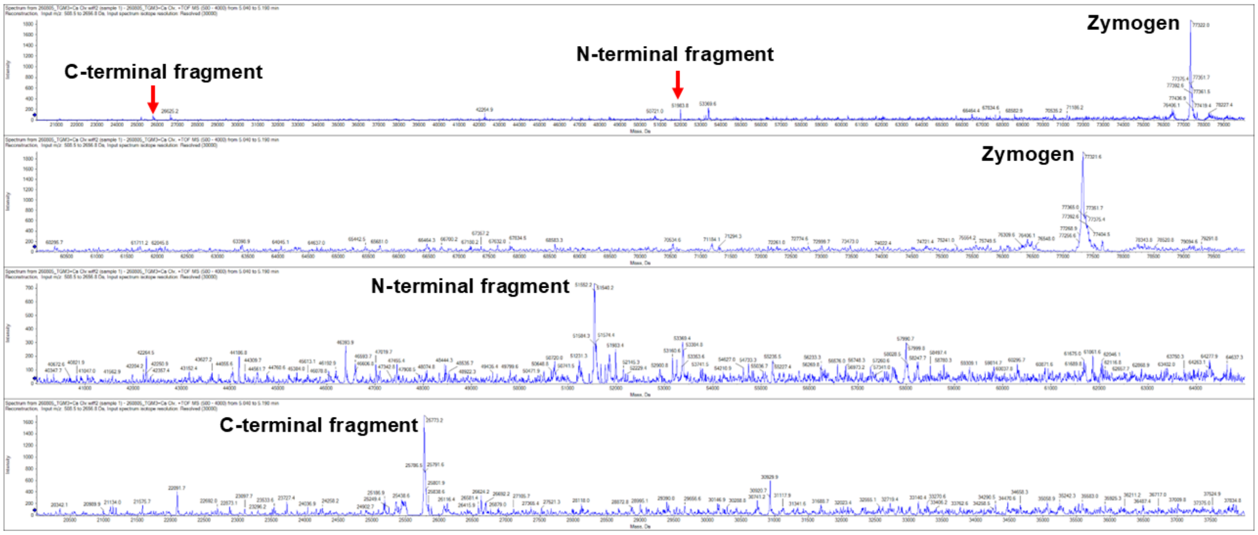


**Figure S3.** High-resolution protein mass spectra of recombinantly purified TG3, indicating its self-cleavage and activation.

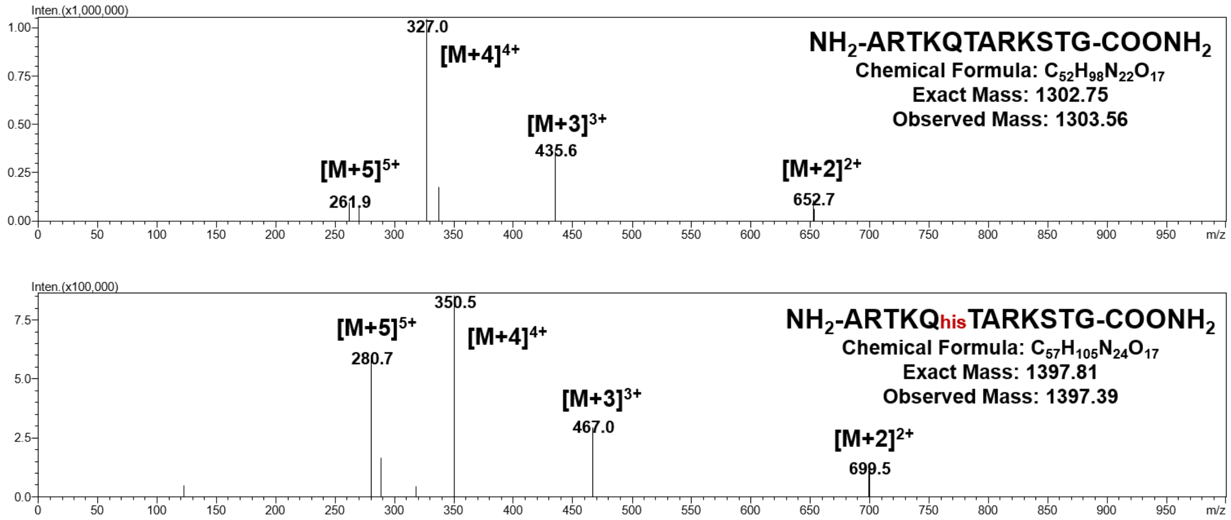


**Figure S4.** Sequences and MS analyses of the substrate peptide H3Q5 (1-12 aa) (top) used for *in vitro* biochemical tests in this study and its corresponding product H3Q5his (1-12 aa) (bottom) converted by TG3.

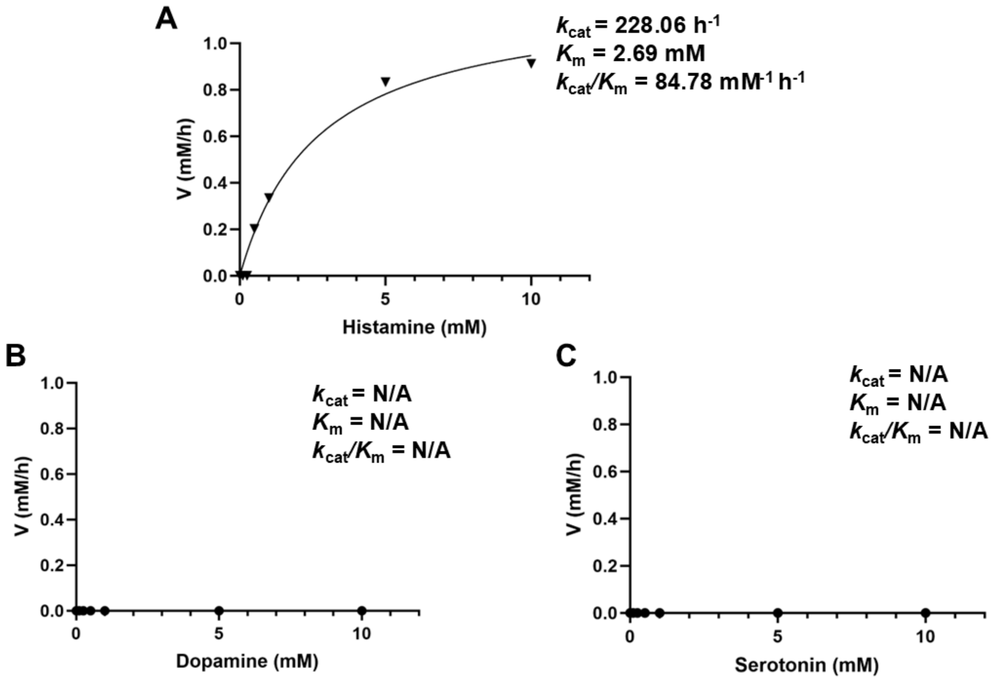


**Figure S5.** Enzyme kinetics analyses of TG3-catalyzed transamidation reactions in the presence of H3Q5 (1-12 aa) peptide and different monoamine substrates, including (**A**) histamine, (**B**) dopamine, or (**C**) serotonin.

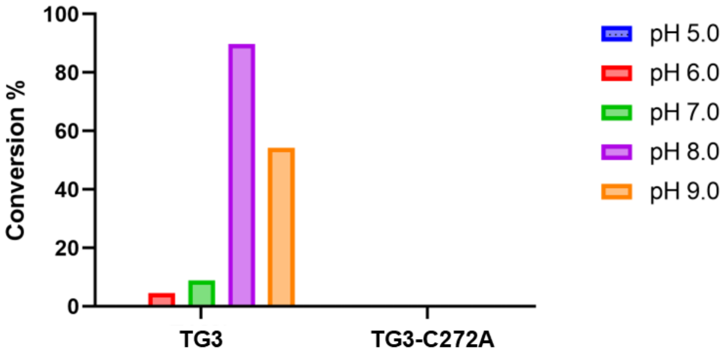


**Figure S6.** pH-Dependence analysis of TG3-catalyzed H3Q5 histaminylation reactions *in vitro*, using the H3Q5 (1-12 aa) peptide as a substrate.

(4) Lee, W.-S.; Al-Ramahi, I.; Jeong, H.-H.; Jang, Y.; Lin, T.; Adamski, C. J.; Lavery, L. A.; Rath, S.; Richman, R.; Bondar, V. V.; Alcala, E.; Revelli, J.-P.; Orr, H. T.; Liu, Z.; Botas, J.; Zoghbi, H. Y. Cross-Species Genetic Screens Identify Transglutaminase 5 as a Regulator of Polyglutamine-Expanded Ataxin-1. *J. Clin. Invest. 132* (9), e156616.

(5) Zhang, N.; Wu, J.; Hossain, F.; Peng, H.; Li, H.; Gibson, C.; Chen, M.; Zhang, H.; Gao, S.; Zheng, X.; Wang, Y.; Zhu, J.; Wang, J. J.; Maze, I.; Zheng, Q. Bioorthogonal Labeling and Enrichment of Histone Monoaminylation Reveal Its Accumulation and Regulatory Function in Cancer Cell Chromatin. *J. Am. Chem. Soc.* **2024**, *146* (24), 16714-16720.

(6) Kumar, S.; Stecher, G.; Suleski, M.; Sanderford, M.; Sharma, S.; Tamura, K. Molecular MEGA12: Evolutionary Genetics Analysis Version 12 for Adaptive and Green Computing. *Mol. Biol. Evol.* **2024**, *41* (12), msae263.
